# A Flexible Framework for Assessing the Cumulative Effects of Offshore Wind Energy Activities and Other Pressures on Aerofauna

**DOI:** 10.1101/2024.08.20.608171

**Authors:** Megan C. Ferguson, Kathryn A. Williams, M. Wing Goodale, Evan M. Adams, Paul Knaga, Katrien Kingdon, Stephanie Avery-Gomm

## Abstract

The offshore wind energy (OSW) industry is pivotal for renewable energy transition and climate resiliency. However, OSW activities may negatively affect aerofauna, contributing to CE from human activities and natural processes. Cumulative effects assessments (CEA) are vital for informed planning and management of OSW activities during regional assessment, site selection, and site evaluation. To reduce impacts on aerofauna, OSW developments should be sited in areas that avoid or minimize cumulative effects (CE). This study presents a cohesive and flexible framework for assessing the CE from OSW activities, other human activities, and natural processes on aerofauna to support decision-making during the initial OSW planning phases. The framework uses a species-based approach, applicable to various receptors, and adapts to available information on ecology, socioeconomics, and pressures. The analytical strategy uses a CE metric to indicate the presence or magnitude of effects from all pressures on receptors. Spatially explicit optimization methods identify OSW site configurations that minimize a CE metric. The framework accommodates alternative pressure scenarios that include foreseeable future human activities and natural processes and can explore the sensitivity of the result to uncertain parameters. Given sufficient spatial information on receptor density, pressure magnitude, and cause-effect pathways, the spatial optimization algorithm can find solutions that minimize population-level impacts from CE. If this ideal standard cannot be achieved due to information gaps, alternative metrics may be used to inform the immediate decision-making process.

## 1.0 Background

The pace at which offshore wind energy (OSW) developments are being built is increasing rapidly, fueled by increasing energy demands and recognition that transitioning to clean, renewable energy is critical to meeting climate targets (Bilgili and Alphan 2022). However, OSW activities also pose risks for wildlife, including aerofauna (i.e., birds and bats, Williams et al. 2024). Mitigation of these impacts follows a near-universal hierarchy: for the priority in which mitigation measures are to be considered and applied: avoidance, minimization, restoration, and compensation (Council on Environmental Quality 2020). Avoidance, achieved through careful siting of OSW developments away from high-risk areas, remains the best available option for mitigating impacts on aerofauna (Gulka et al., 2024 preprint).

Decisions regarding the future of the OSW industry will differentially impact people, wildlife populations, and ecosystems. The effects of OSW on wildlife should be considered in the context of other anthropogenic and environmental pressures. The term “cumulative effects” generally refers to effects that may be individually minor, but collectively significant. We define cumulative effects (CE) as the combined effects of human activities and natural processes on wildlife across space and time (i.e., past, present, and reasonably foreseeable future). The term ‘reasonably foreseeable’ refers to something that is likely or expected to occur. A cumulative effects assessment (CEA) is a systematic process of identifying, analyzing, and evaluating CE on a receptor, for the purpose of informing planning and management. Ideally, a CEA should be conducted at the regional scale early in the planning process so that information is available to guide planning decisions, including where to site OSW licensing areas. The amount and types of information available to include in a CEA varies by species, location, and time.

In Canada, the OSW industry is emerging. Federal and provincial governments are designing regional strategies for managing future activities related to OSW developments in Nova Scotia and Newfoundland and Labrador (i.e., Regional Assessments; Committee for the Regional Assessment of Offshore Wind Development in Nova Scotia 2024; Committee for the Regional Assessment of Offshore Wind Development in Newfoundland and Labrador 2024). In particular, under the authority of the Impact Assessment Act of 2019 (IAA), the Regional Assessment Committees are tasked with identifying and considering the potential positive and adverse effects, including CE, of future OSW development activities in the two regional study areas, as well as “potential interactions between the effects of future offshore wind development activities and those of other existing and future physical activities, including the potential for resulting cumulative effects”. Currently, Canada does not have a cohesive framework for conducting regional CEAs, which presents an opportunity to develop such a framework.

To support the sustainability of the emerging OSW industry in Canada and worldwide, this paper presents a framework for assessing the CE of OSW activities, other human activities, and natural processes on aerofauna at a regional scale. First, we synthesize best practices, including fundamental CEA concepts and basic steps for a species-centric CEA that could be applied to a variety of wildlife receptors (i.e., populations, species, or valued components). Then we present the analytical strategy, providing both a mathematical and a verbal description of the CE metrics, and a verbal description of the spatial optimization algorithm. While conceptually similar to Halpern et al. (2008), our approach extends its utility by allowing the variables related to receptors and pressures that are required to compute the CE metric to vary depending on available information, enhancing the flexibility and applicability across different receptors and pressures. We clearly show how a number of superficially different approaches to conducting a CEA really fit under a single umbrella, and we explain how analyses based on different information types can complement each other in a single analysis. This framework is suitable for delineating areas where OSW developments are most likely to minimize CE. A glossary is provided to facilitate a common understanding (Appendix A).

## 2.0 Phases of OSW Planning

A CEA may be used to inform decision-making during three distinct phases of OSW planning: (i) regional assessment and region delineation (i.e., delineating regions within which OSW activities may occur); (ii) site selection for OSW development activities; and (iii) OSW site evaluation (Figure 1). These planning phases ideally should occur in the order listed.

**Figure 1.**
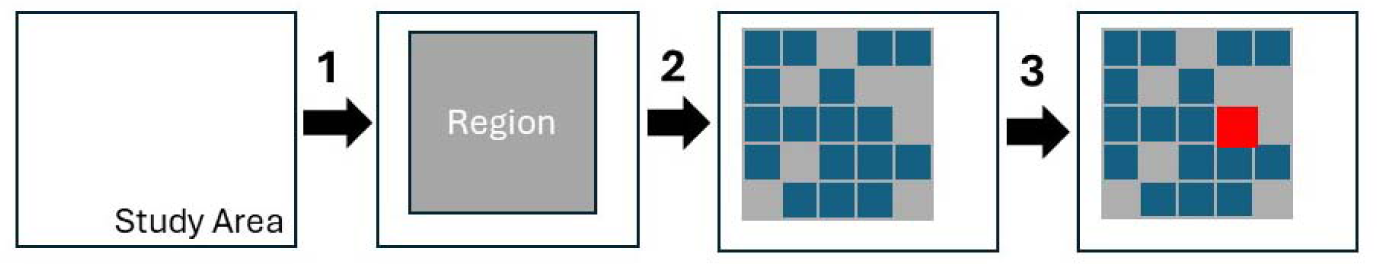
Schematic representation of three phases of offshore wind planning: 1) regional assessment and region delineation; 2) site selection; and 3) site evaluation. The study area is the largest geographic area considered for future offshore wind energy development activities; it may coincide with federal or provincial jurisdictional boundaries. Regional boundaries represent the first level of refinement in the spatial planning process (e.g., “Preliminary Offshore Wind Licencing areas” or “Potential Future Development Areas”; Committee for the Regional Assessment of Offshore Wind Development in Newfoundland and Labrador (2024); Committee for the Regional Assessment of Offshore Wind Development in Nova Scotia (2024)). The small blue squares represent individual project areas (i.e., licencing or lease areas that developers may bid on), which are delineated within the regional boundaries during site selection. During site evaluation, a cumulative effects analysis is conducted on one specific site, highlighted in red in the figure.

Phase 1, **Regional assessment and region delineation**, describe the processes used to analyze and evaluate OSW development scenarios, with the goal of informing and improving future planning, licencing, and impact assessment processes^1^. The identification of “Preliminary Offshore Wind Licencing areas” and “Potential Future Development Areas” by the Committees for the Regional Assessments of offshore wind development in Newfoundland and Labrador, and Nova Scotia (respectively) are examples of this phase (Figure 2; Committee for the Regional Assessment of Offshore Wind Development in Newfoundland and Labrador 2024; Committee for the Regional Assessment of Offshore Wind Development in Nova Scotia 2024).

**Figure 2.**
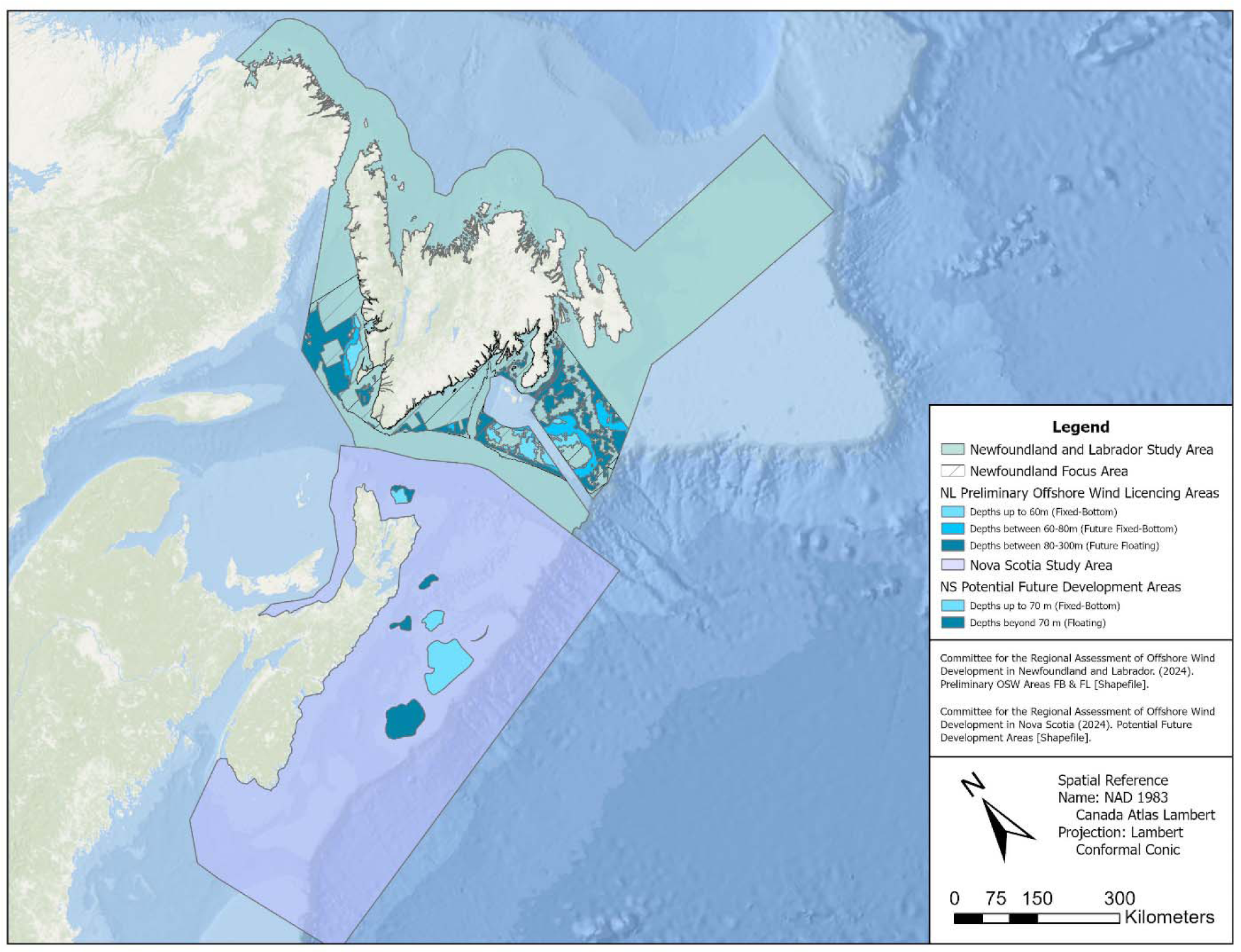
Preliminary Offshore Wind Licencing Areas in Newfoundland and Labrador (Committee for the Regional Assessment of Offshore Wind Development in Newfoundland and Labrador 2024) and Potential Future Development Areas for offshore wind in Nova Scotia (Committee for the Regional Assessment of Offshore Wind Development in Nova Scotia 2024).

Phase 2, **site selection**, occurs within pre-defined regional boundaries. This is the process of identifying specific boundaries for project areas that developers may bid on. Regional assessments, region delineation, and site selection encompass large geographic areas.

Phase 3, **site evaluation**, contrasts with phases 1 and 2 because it involves assessing the potential CE or impacts from one specific OSW project at a particular location, considered alongside a range of other OSW projects in the region or other types of activities influencing the same receptors (i.e., ecosystems or wildlife populations).

Our framework was designed to identify the best strategies for delineating boundaries of future potential OSW developments within a larger geographic area, with the aim of minimizing Ce of human activities and natural processes on wildlife. Therefore, the framework can support the identification of regional boundaries within a larger study area during the **regional assessment and region delineation** phase, and the delineation of individual project boundaries within a prespecified region during the **site selection** phase. Following the regional assessment process and leasing of offshore areas for wind energy development, site-specific CE evaluations (i.e., phase 3 **site evaluations**) should be conducted. The products developed during phases 1 and 2 may inform the site evaluation phase. For example, information about species and pressures that is compiled during the first two phases may be incorporated into future site evaluations.

Additionally, the results of regional assessment, region delineation, and site selection could provide regional context to inform the design and interpretation of site evaluations.

## 3.0 Fundamental CEA Concepts

In this section we address four fundamental concepts related to CEAs. We begin by defining the scope of a CEA (*Section 3.1*). The scope comprises several variables that are used to define the extent of the CEA. Defining the scope is critical for constructing a problem that can be solved reasonably and for communicating the utility of the CEA results. A clear statement of the management objectives is essential for refining the scope of the CEA. In *Section 3.2*, we provide examples of the range of management objectives that CEAs have addressed. In *Section 3.3* on typology, we identify four general types of CEA frameworks and provide our definition for a “species-based approach”. Lastly, we list and describe the fourteen basic steps involved in conducting a species-based CEA (*Section 3.4*).

### 3.1 Scope

Conceptually, an environmental CEA is all-encompassing: it includes all effects from all anthropogenic activities and natural processes on all environmental components, with no spatial or temporal constraints. However, in practice, every pressure, interaction, and effect cannot be understood and analyzed. Without narrowing the scope of a CEA down from “all-encompassing” to a justifiable and manageable subset of possibilities, the CEA is intractable (Goodale and Milman 2016; Masden et al. 2010; Adams et al. 2023; Brignon et al. 2022).

In a CEA, the term “scope” refers to the envelope or boundaries defining: the (i) spatial and temporal constraints of the analysis; (ii) receptors (e.g., populations, species, valued components, or ecosystems) included in the analysis; (iii) anthropogenic activities (sources) and specific pressures posed by those activities that are assessed; and (iv) future development scenarios that are assessed (e.g., total area occupied by OSW development or total energy produced).

### 3.2 Management Objectives

The management objectives, and hence the objectives for a CEA, will predominantly guide the definition of the CEA’s scope variables and selection of analytical methods. A non-comprehensive list of the types of management objectives that CEAs in Europe, the United States, and Canada have addressed, either alone or in combination, include:

1. identifying regional boundaries within which development can occur (i.e., phase 1 defined in *Section 2: Phases of OSW Planning*);
2. site selection (i.e., phase 2 defined in *Section 2: Phases of OSW Planning*);
3. assessing and evaluating the risks to receptors from different anthropogenic sources;
4. identifying actionable measures to avoid, minimize, or compensate for cumulative adverse effects to wildlife from human activities; and
5. evaluating estimated CE metrics with respect to pre-established criteria (i.e., thresholds, benchmarks).

Our CEA framework can assist with each of these objectives, given appropriate specification of the scope variables (*Section 3.1: Scope)* and alternative scenarios (*Section 4.7: Scenario-building and Sensitivity Testing*).

### 3.3 Typology

Murray et al. (2020) categorize CEAs into four typologies: activity-based, stressor-based, species- or habitat-based, and area-based frameworks. Our CEA framework applies a population- or species-based approach, wherein the ecology of the receptor species is the primary factor used to constrain the spatial, temporal, and activity/pressure scope variables (E. Willsteed et al. 2017; Murray, Hannah, and Locke 2020). In the text that follows, we use the term “species” for simplicity. For cases in which population structure is known or can be inferred, it is best from an ecological perspective to base the analysis on the population rather than the entire species. A population comprises individuals that can potentially interbreed, which is important for effective conservation and management. However, for cases in which relevant laws or regulations apply to species, it may be more appropriate to base the analysis on the species.

### 3.4 Basic Steps for a Species-based CEA

Regardless of the OSW planning phase for which a CEA is prepared (regional assessment and delineation; site selection; or site evaluation), we propose a species-based CEA that includes the following steps:

1. Explicitly define the objectives of the CEA (Stelzenmüller et al. 2018; E. A. Willsteed et al. 2018). The CEA objectives should align with the designated management objectives.
2. Identify the specific geographic boundaries of the OSW area(s) of interest (i.e., the region or the OSW development project).
3. Inventory the species that occur within the geographic boundaries for the area of interest that were specified in Step 2. This should include species that occur in the area only occasionally (e.g., migratory or other species with spatiotemporal variability in their distribution) and seasonal and permanent residents. The initial species inventory should be comprehensive, including species that rarely occur in the area or whose distribution is uncertain but may overlap the area at some point in time (Brignon et al. 2022). The inventory should be based on the best available information and may be supported by expert judgement where information is scarce.
4. Identify all known potential cause-effect pathways that link the species to known OSW pressures. This step can be based on expert knowledge and a review of the available literature. It is important to define which development phases (i.e., construction, operation, or decommissioning) and the type of OSW technology (e.g., fixed foundation or floating platform) will be included in the scope. At this step, the list of pathways should be comprehensive; the particular pressures included in the analysis will be refined in Step 9.
5. Select the receptor species that will be included in the CEA from the inventory of species identified in Step 3. Factors to consider in selecting the receptor species are listed below (*Section 4.1: Selection of Receptor Species*).
6. Refine the spatial scope. For a species-based assessment, the spatial extent of the regional or project-level OSW development activities that prompted the CEA (identified in Step 2) is likely to be smaller than that of a given receptor species’ range. A comprehensive species-based CEA would investigate all pressures affecting the species throughout its range. As discussed further below, a hierarchical approach can be used to define the spatial scope of the analysis, allowing differences in the types and resolution of information that are used to assess effects within the OSW activity area compared to other parts of the species’ range.
7. Inventory the other primary sources (i.e., non-OSW activities and natural processes) and associated pressures occurring anywhere in the ranges of the selected receptor species.
8. Identify all known potential cause-effect pathways that link the species and the non-OSW pressures listed in Step 7. This step can be based on expert knowledge and a review of the available literature. See *Section 4.2: Identifying Cause-effect Pathways* for more details.
9. Select the OSW and non-OSW pressures that will be included in the CEA. Factors to consider include, inter alia, understanding of the cause-effect pathway linking the pressure to the receptor species, sensitivity of the receptor species to the pressure, and availability of information needed to incorporate the pathway into the CEA. If high-risk pathways are identified that have insufficient information to formally analyze, they can be added to a list to help prioritize future research.
10. Define the temporal scope of analysis. This requires specifying baselines for the receptor species and how far into the future the analysis will predict. Factors to consider in determining baseline conditions are discussed below (*Section 4.3: Baselines*).
11. Define the metric(s) (i.e., variable(s) or parameter(s)) that will be used to measure CE and the criteria (i.e., benchmarks or thresholds) that will be used to evaluate the significance of the resulting effects (Stelzenmüller et al. 2020). The decision-making criteria should be chosen with objective input from the wildlife managers, environmental and energy regulators, and experts in the different types of criteria that have been used in other cases. The advantages and disadvantages of existing criteria should be weighed relative to the management objectives and relevant regulations. Ideally, candidate criteria should undergo simulation testing to evaluate whether they are likely to provide the necessary information to the decision-makers. The CEA metric(s) should provide the information required to apply the decision-making criteria. The CEA metric(s) and criteria should be standardized across all assessments to allow the results of different assessments to be compared and to facilitate consistency in the decision-making process. See *Section 5.2: CEA Metrics* for more details.
12. Define a set of scenarios with reasonably foreseeable changes in OSW pressures, non-OSW anthropogenic pressures, or natural processes. This step will benefit from diverse input (i.e., from energy and environmental regulators, developers, scientists, Indigenous peoples, and stakeholders; Duinker and Greig 2021; 2007). For OSW, scenarios may involve different types of technology or different buildout goals that are defined in terms of total energy produced or total area covered by OSW activities. Multiple types of human activities and natural processes can be included in scenarios. See *Section 4.7: Scenario-building and Sensitivity Testing* for more details.
13. Estimate the values of the CE metric(s) and associated uncertainty for each scenario (e.g., Searle et al. 2023; Stelzenmüller et al. 2020). For more details see *Section 4: Additional Considerations*and *Section 5.3: Spatial Optimization Algorithm*.
14. Compare the estimated CE metric(s) to the evaluation criteria to inform spatial planning and CE management decisions by decision-makers.

## 4.0 Additional Considerations

In this section, we discuss seven additional factors that influence CEA results: receptor species (*Section 4.1*); cause-effect pathways (*Section 4.2*); definition of baselines (*Section 4.3)*; spatial variability in marine habitats (*Section 4.4*); sources of uncertainty and methods for estimating uncertainty in CEA results (*Section 4.5)*; data availability, representativeness, and quality (*Section 4.6*); and scenario uncertainty (*Section 4.7*).

### 4.1 Selection of Receptor Species

The receptor species and human activities selected for inclusion in a CEA depend on the OSW planning phase (i.e., regional assessment and region delineation, site selection, or site evaluation). They also depend on the spatial extent of the OSW activities for which the CEA is conducted, because the prioritization of a species or activity may vary with the geographic location or size of the analysis area. There are many factors to consider when selecting receptor species (Masden et al. 2010; Tulloch et al. 2024; Lerner 2018; Popper et al. 2022; Regional Synthesis Workgroup of the Environmental Technical Working Group 2023), including:

- conservation status, based on either official designation (provincial, national, international) or inferred vulnerability to predicted future ecological or anthropogenic changes;
- number of individuals or proportion of the population that uses the area;
- age, age class, or sex of individuals that use the area;
- activity/activities undertaken in the area (e.g., feeding, migrating, breeding, molting, nesting, or rearing young);
- known or suspected vulnerability to OSW activities in the area;
- life history parameters, such as long lifespan and low reproductive output, that would make the species particularly sensitive to disturbance;
- importance to Indigenous peoples or stakeholders;
- ecosystem functioning and trophic interactions, because the presence, absence, or abundance of certain species may have considerable influence on the ecosystem, or the species may represent a broader collection of taxa, such as a guild, community, or ecosystem; and
- availability of information needed to conduct the CEA.

If a species ranks high in priority based on these factors, but insufficient information exists to estimate any of the CEA metrics described in *Section 5.0 Analytical Strategy*, the species can be added to a list to help prioritize future research.

### 4.2 Identifying Cause-effect Pathways

Pathways of effects modelling (PoE) is a useful tool to systematically identify cause-effect pathways between human activities, the associated stressors to ecological components, and the expected effects (Government of Canada 2012; Knights, Koss, and Robinson 2013). PoE development involves defining measurable endpoints (e.g., habitat distribution, availability) that vary based on management objectives (Government of Canada 2012). Mapping PoE includes reviewing and synthesizing existing knowledge on pressures, mechanisms, and potential effects. PoE models provide simplified visual representations of complex interactions, which allow opportunities to further identify more complex CE and possible mitigations (Clarke Murray, Mach, and Martone 2014; Isaacman and Daborn 2011; Knights, Koss, and Robinson 2013). By developing PoEs, all known potential effects or impacts of pressures on receptors can be considered and addressed, ultimately contributing to more comprehensive and accurate CEAs.

### 4.3 Baselines

The baseline sets the standard against which comparisons are made. The baseline for a species is defined by a specific period and abundance or effective population size. Factors to consider when setting baselines may include the following:

- formally designated conservation targets for individual populations or species;
- best available information on abundance immediately prior to development (but see “shifting baseline syndrome,” below);
- estimated trends in population abundance over time;
- maximum historical abundance estimate;
- Indigenous knowledge;
- uncertainty in existing abundance estimates;
- spatiotemporal variability in the distribution of the receptor relative to the distribution of effort used to estimate abundance, because it is typically more difficult to estimate abundance for receptors whose distributions are highly variable;
- history of known mortality, such as from directed harvest, bycatch, diseases, oil spills;
- estimates of critical parameters, such as carrying capacity or maximum net productivity level, from mathematical models of population dynamics; and
- changes in environmental conditions within the area over time.

The aforementioned list of factors includes information compiled from Masden et al. (2010), Adams et al. (2023), Bangay et al. (2020), Goodale and Milman (2016), Warwick-Evans et al. (2018), Kelsey et al. (2018), BOEM (2020; 2024), Jongbloed et al. (2023), Peschko et al. (2024), Halpern et al. (2008), Goodale et al. (2019), Rijkswaterstaat (2022), Potiek et al. (2022), Tulloch et al. (2024), Robinson Willmott et al. (2013), and Layton-Matthews et al. (2023).

Sources of this type of information for wildlife in Canada include Committee on the Status of Endangered Wildlife in Canada (COSEWIC) status reports^2^, Species at Risk Act recovery strategies^3^, and conservation strategy reports for Bird Conservation Regions and Marine Biogeographic Units (Environment Canada 2013).

Care must be taken to avoid the “shifting baseline syndrome” (Pauly 1995), which occurs when a population abundance decreases over time, yet the time series of abundance is ignored when setting the baseline. As a result, the baseline reflects only the most recent abundance estimate, leading to a gradual decline in the baseline and, hence, management targets that are insufficient for conservation or management purposes.

### 4.4 Spatial Variability in Marine Habitats

Depending on how a species uses the marine environment, CE may be compounded or mitigated by clustering all OSW activities in a confined subarea; analogously, broadly dispersing activities across a vast area might compound or reduce CE for certain species (Masden et al. 2010; Goodale, Milman, and Griffin 2019). The best way to avoid negative effects of OSW activities on a receptor species is to locate OSW activities outside of the receptor’s range. If that option is not feasible, several factors related to marine habitats should be considered when applying the CEA framework to effectively understand and minimize CE. The most important factor is that marine habitats are spatially heterogeneous; this is true for aerofauna and for taxa that spend most or all their lives underwater. For example, geographic, topographic, and bathymetric features such as seamounts, canyons, shelf breaks, estuaries, points, and bays may enhance biological productivity or physical aggregation and retention of resources, which may result in high densities of upper trophic level species. Bathymetry or shoreline topography may create areas that are advantageous for breeding or rearing young. The strength and direction of air currents and winds affect aerofauna flight efficiency; these atmospheric phenomena exhibit spatiotemporal patterns and variability that interact with oceanographic phenomena to link ecosystems in time and three-dimensional space, resulting in spatiotemporal variability in aerofauna density.

### 4.5 Uncertainty

When considering the results of a CEA and comparing the estimated CE metric(s) to the evaluation criteria, decision-makers need to know how much confidence to put in the results. Confidence and uncertainty are inversely related: we are less confident in things that are highly uncertain, and more confident in things that have low uncertainty.

In the field of CE, “uncertainty” can be defined in relatively broad terms as “the state of deficiency of information related to understanding or knowledge of an event, its consequences or likelihood” (Stelzenmüller et al. 2018). Uncertainty in a CEA may be classified into four categories:

- **Observation uncertainty** could be due to random errors (i.e., precision) or systematic discrepancies in magnitude or direction between data and reality (i.e., bias) (Stelzenmüller et al. 2020).
- **Process uncertainty** in ecology refers to incomplete knowledge of relationships among natural phenomena, such as interactions among species and how species relate to their environment (Cressie et al. 2009). Ecological process uncertainty also includes incomplete knowledge of the relationships between natural phenomena and human activities (e.g., cause-effect pathways, *Section 4.2*).
- **Statistical uncertainty** refers to uncertainty in the assumptions used in statistical models; examples include model structure or parameter estimates (Stelzenmüller et al. 2020).
- **Pressure scenario uncertainty** arises due to imperfect knowledge about the range and intensity of future human activities (*Section 4.7*: *Scenario-building and sensitivity testing*).

For further details on uncertainty in CEAs, see Stelzenmüeller et al. (2020).

The sources and magnitudes of uncertainty, and the methods used to account for uncertainty, will affect the results of a CEA. Some CEAs ignore uncertainty; others use qualitative (e.g., ranks) (Robinson Willmott, Forcey, and Kent 2013; Kelsey et al. 2018; Potiek et al. 2022) or quantitative methods (e.g., Monte Carlo simulations, Potiek et al. (2022) and Soudijn et al. (2022); Bayesian networks, Tulloch et al. (2024)) to try to estimate and propagate uncertainty from the input variables to the final CE metric(s). When unable to explicitly account for uncertainty in an analysis used to estimate seabird mortality due to collisions with wind turbines in the North Sea, Potiek et al. (2022) relied on the precautionary principle and assumed the “worst case scenario”, which they defined as the scenario that would result in the highest mortality. The degree to which a CE metric responds to uncertainty may be investigated using statistical methods such as sensitivity testing, which is discussed further in *Section 4.7: Scenario-building and Sensitivity Testing*.

### 4.6 Data Availability, Representativeness, and Quality

Observation uncertainty is a function of the amount of data available (often referred to as sample size) and how well the data (the sample) represents the underlying truth. Observation uncertainty can be quantified using precision and bias.

Precision is known by several terms, including random error, variability, random variation, and noise. Precision has no preferred direction, and increasing sample size should effectively increase precision. Quantifying the size of a sample of biological data is not always straightforward. The following are guidelines on criteria for evaluating sample sizes for a variety of types of ecological data, modified from Harrison et al. (2023):

- **Bio-logging**: number and type of tags deployed in different age and sex classes or populations at particular locations and times; tag longevity (i.e., the length of the time series from each tag); and the number of years across which the tags were deployed on a particular population or species;
- **Opportunistic visual observations**: number of observations and the temporal and spatial extent and resolution of the observation effort;
- **Photo-ID**: number of individuals identified; study duration (in years); spatial and temporal extent of sampling; representativeness of the sample (e.g., age class, sex, proportion of the population with identifiable markings); number of resightings of individuals; and the maximum number of years a single individual has been identified in an area;
- **Line-transect survey**: number of surveys in the time series; number of observations; number and length of the transects; effective strip width; time lag between surveys; and the temporal and spatial extent and resolution (i.e., spacing between transects) of each survey; and
- **Passive acoustic monitoring (PAM)**: number and location of acoustic recorders; spatial and temporal extent of recordings; sample frequency; and number of signals (i.e., calls, whistles, clicks, songs, etc.) of the specific species detected.

Bias has a net direction and magnitude regardless of sample size and is therefore a measure of inaccuracy. In general, bias can arise during the sample design, data collection, or analytical steps when the underlying assumptions are not valid for the case study. A biased dataset may result from systematically preferring (or avoiding) objects with particular characteristics during sampling, resulting in a skewed or unrepresentative sample of reality. Bias may inadvertently arise during data collection when inclement weather precludes sampling the entire survey design. Other examples of data that may be considered biased for a particular case study include data that were collected in the following ways: on species other than the receptor species in the CEA; in a different place than the area of interest to the CEA; during a different period (e.g., month, season, or year) than the period of interest to the CEA. Bias may also result from collecting data using a tool that is poorly calibrated so that the measurements are all either too low or too high. Bias can also arise during analysis due to the estimator chosen or the methods used to analyze the data. When sampling bias is known to exist and methods are implemented to estimate and correct for the bias during analysis, the bias does not propagate to the results of the analysis.

Data quality is related to precision and bias and is case specific. For example, Potiek et al. (2022) modeled the demographics of seabirds in the Dutch part of the North Sea. To help represent the uncertainty in their analysis, they scored the data sources used to derive the input parameters needed for the population modeling. The scores were ordinal (0, 1, or 2) based on data quality and representativeness. They scored data quality based on the number of years in the data, the number of individuals in the data, and whether uncertainty was reported. They scored representativeness based on data recency, how representative the data were to their study area, and how representative the data were to the current trends in seabird populations in their study area.

### 4.7 Scenario-building and Sensitivity Testing

One method of examining the sensitivity of CEA results to changes in parameters or model structure that are not known with certainty is to implement the CEA independently in multiple scenarios. For example, the scale and effects of OSW, non-OSW human activities, and natural processes (e.g., climate change) on wildlife may be poorly known in some instances. In these cases, pressure scenarios should define a range of pressures that could arise from the sources included in the CEA (*Section 3.4: Basic Steps for a Species-based CEA, Step 12*). In general, each scenario uses scenario-specific parameters to address one or more unknowns, such as: total OSW buildout (e.g., measured in terms of total energy produced or area covered by OSW projects); effects of other anthropogenic activities on the receptor, taking into consideration existing and hypothesized future management decisions; climate change effects; current receptor abundance; life history parameters; and cause-effect pathways. Comparing CEA results from multiple scenarios can help to distinguish between parameters that heavily influence CEA results (i.e., a small change in the parameter value generates a large change in the CE metric) and parameters that have only weak influence on the results (i.e., a large change in the parameter value generates a negligible or small change in the CE metric). Understanding the sensitivity of the CEA results to different parameters can help prioritize future research, with the ultimate goal of enhancing the certainty of future CEAs and the effectiveness of future management decisions.

The conclusions that can be drawn from the analysis of multiple scenarios are only as good as the suite of scenarios that are investigated. The future is unpredictable. It is acceptable if the actual progression of events does not align with any of the scenarios analyzed because sensitivities can still be investigated using a well-designed collection of scenarios. Input from Indigenous peoples, developers, scientists, stakeholders, resource managers, and environmental and energy regulators may be used to define the set of scenarios to analyze. We present general guidelines for scenario-building, and then provide an example of how those general guidelines may be applied specifically to designing pressure scenarios for CEAs focused on OSW energy activities. The general guidelines, which were based on those provided in Duinker and Greig (2007), are as follows:

- The set of scenarios should include sharp contrasts in the unknown parameter(s);
- The set of scenarios should comprehensively span the range of likely values of the unknown parameter(s);
- Each scenario should be rooted in the present (e.g., climate change forecasts should be empirically grounded in the present to provide confidence that they begin in the right place), plausible (not impossible), and internally consistent;
- Our collective ability to judge probabilities of outcomes is poor. Therefore, avoid trying to create a “most likely” scenario. This applies also to creating three scenario clusters with some notion of “high”, “medium” and “low”, and to classifying the likelihood of future events into “almost certain,” “reasonably foreseeable,” and “hypothetical”; and
- The total number of scenarios that are evaluated should represent a balance between parsimony (i.e., few scenarios) to make the analytical task tractable, and comprehensiveness (i.e., many scenarios) to disentangle interactions among parameters for realism.

The first two general guidelines can be adapted to the specific task of designing pressure scenarios for a CEA focused on OSW activities as follows:

- The set of scenarios should include sharp contrasts in alternative futures, as defined by factors with substantial uncertainty such as: OSW buildout and technology; climate change; management decisions that affect the receptor species (e.g., management actions that mitigate the effects of bycatch, light pollution, contaminants, noise, competition for prey, disease);
- The set of scenarios should comprehensively span the range of potential future OSW buildout and technologies, and all key pressures with the potential to measurably affect the receptors, either individually or cumulatively. Pressure sources may be primarily anthropogenic (e.g., direct effects from fisheries, shipping, tourism, military), environmental (e.g., inter- or intra-specific competition for resources, disease) or a combination of both (e.g., climate change).

## 5.0 Analytical Strategy

### 5.1 Overview

Briefly, our analytical strategy uses spatial optimization methods that aim to minimize the value of a CE metric via the spatial allocation of individual polygons within a larger geographic area. Thus, it can be used both when the CEA is intended to inform phase 1 (i.e., regional assessment and region delineation) or phase 2 (site selection). The combination of possible polygons that may be selected to form a valid solution to the optimization problem may be constrained by factors such as minimum or maximum OSW site size, total area of all OSW sites within the region, total energy produced in the region, or avoidance of areas of concern (e.g., based on pre-defined ecological, social, economic, or logistical factors).

Given sufficient spatially explicit information about the density of the selected receptors, magnitude of the selected pressures, and cause-effect pathways linking the receptors and pressures, we can program the spatial optimization algorithm to find solutions that minimize population-level impacts to a receptor or a community from the CE of all pressures. If this ideal standard cannot be achieved due to information gaps, alternative metrics presented below may be used to inform the immediate decision-making process. The list of information gaps may help guide future research efforts.

The CE metric is defined as the product of a spatially explicit species variable and a spatially explicit pressure variable, summed across all species and pressure combinations. Although our approach is conceptually similar to that of Halpern et al. (2008), we extend its utility by allowing the specific variables used to compute the CE metric to vary depending on available information. This type of flexible yet cohesive approach allows for standardization of metrics and criteria across all assessments, enabling the results from different assessments to be compared and facilitating consistency in the decision-making process.

### 5.2 CEA Metrics

Five structurally similar CE metrics can be produced from the species and pressure variables, dependent upon data availability. **Figure 3** shows a schematic representation of how different combinations of species and pressure data can be incorporated into CE metrics that are structurally similar, but that represent diverse types of information about the CE across species, pressures, space, and time.

**Figure 3.**
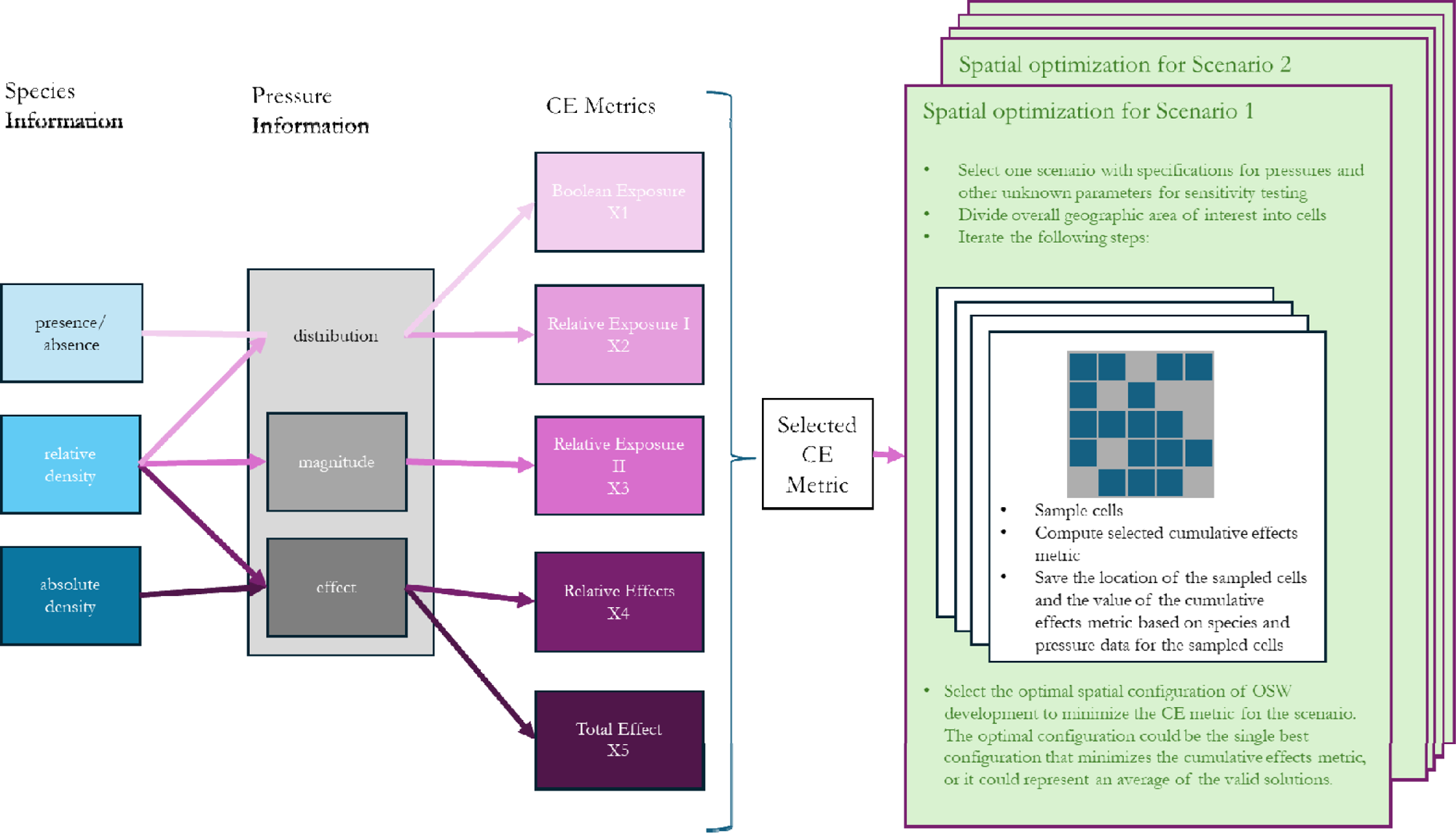
Schematic representation of the species-centric cumulative effects assessment (CEA) framework. Selection of the cumulative effects (CE) metric is influenced by the types and quality of spatial information available on the species and pressures that are included in the pressure scope. In the figure, higher quality information is indicated with darker shading. If spatial information does not exist on species or pressures, cumulative effects assessments using this framework are not possible. Following the selection of the CE metric, scenario-specific parameters are defined for sensitivity testing, and the spatial optimization process is run iteratively. The output for each scenario is the optimal spatial configuration of OSW sites that will minimize the CE metric, or an average of valid solutions.

The variable for species *s* at location *h* may be one of three types: binary presence/presumed absence or presence/absence, *I_s,h_*; relative density, a potentially biased estimate of the number of animals per unit area, 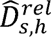; or absolute density, an unbiased estimate of the number of animals per unit area, 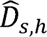.

The variable for pressure *p* at location *h* may also be one of three types: binary presence/presumed absence or presence/absence, *I_p,h_*; estimated pressure magnitude,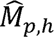; or estimated pressure effect on an individual of a given species, 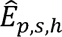. The effect (i.e., proximate response of an individual, such as a change in behavior or diet) that a pressure has on an individual likely depends on the species’ sensitivity to the pressure and on the magnitude of the pressure.

Broadly, the information captured in these five CE metrics increases from the **Boolean exposure** (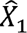) metric to the **total effect** (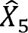) metric, resulting in a concurrent increase in the utility of metrics.

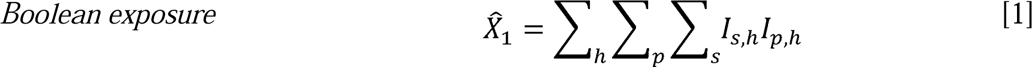

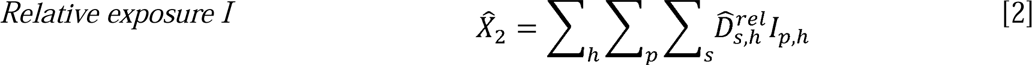

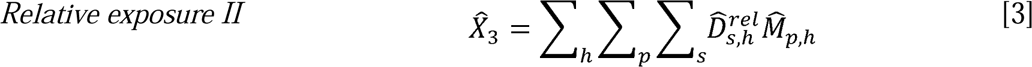

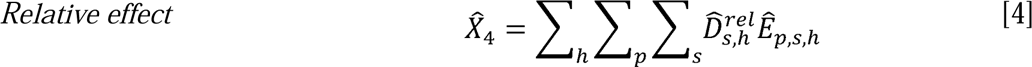

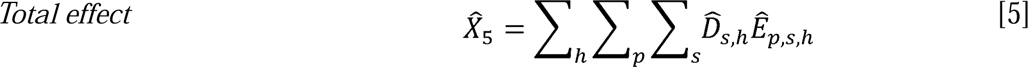

First, we define each of the metrics, then we discuss how more than one metric could be combined within a single CEA.

1. The **Boolean exposure** (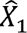) metric represents the sum across all receptor species, pressures, and locations of a spatially explicit presence/absence metric. The 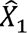 metric can be used only to account for spatial overlap between species and pressures.
2. The **relative exposure I** (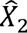) metric provides information about spatial heterogeneity in species’ relative density across the region, but it considers only the overlap of the species and pressures - not the magnitude of the pressures or the pressures’ effects on the species. Furthermore, because 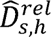 represents only relative density, metric 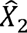 should be applied and interpreted with caution due to the potentially unequal and unknown weighting of each species.
3. The **relative exposure II** (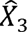) metric also relies on relative species density, but it incorporates information about the spatial heterogeneity across the region in the magnitude of each pressure. Metric 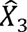 should be interpreted with caution because it is based on a relative species variable that may have biases that are not consistent across all species and on a pressure variable (magnitude) that might not relate consistently across all species to the pressure’s effect on an individual (e.g., if species differ in their sensitivities or proximate responses to the pressure).

Metrics 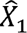, 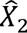, and 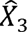 could be converted to vulnerability metrics if information about sensitivity is known for each combination of species and pressure. For example, any one of these three CE metrics could be multiplied by a species-specific sensitivity metric. The resulting spatially explicit vulnerability metric could be used to evaluate the risks from alternative management scenarios, or it could be used in the spatial optimization algorithm defined below to identify strategies for minimizing effects to vulnerable species.

4. The **relative effect** (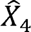) metric provides information on vulnerability or relative effects by weighting the relative density of each species according to each pressure’s effect on the species. However, 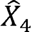 cannot address the absolute number of individuals of each species that are affected because it incorporates species-specific estimates of relative density, which may be biased, and the biases may not be consistent across all species.
5. The metric with the most utility is the **total effect** (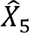) metric because it is an unbiased estimate of the total effect of each pressure on each species. In the case of only a single species, if the effect of a pressure on an individual is death, then 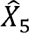 represents the total number of individuals that die from the pressures included in the scope of the analysis. Alternatively, if the effect of a pressure is reduced reproductive success, then 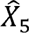 could represent a reduction in the size of the breeding population, number of offspring produced, or number of offspring surviving to reproductive age.

Metric 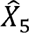 could be used in a population viability analysis (PVA) to infer population-level impacts, such as the magnitude of the change in abundance over time, variation in population growth rates, or the probability that abundance will change within certain parameters over a specified period. Additionally, 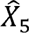 could be multiplied by an estimate of population vulnerability (e.g., conservation status) to inform risk-based management decisions.

Note that it is possible to multiply absolute species density by either a binary pressure variable (*I_p,h_*) or the magnitude of the pressure (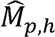). However, we do not include either of those metrics as distinct options because the result of either operation would be essentially another relative exposure metric. Although the number of individuals being exposed would be assumed to be known without bias (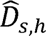), there would be insufficient information about the pathway of the pressure’s effect on individuals (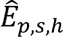) to estimate the total effect of the pressure to the species.

### 5.3 Spatial Optimization Algorithm

The generalized spatial optimization algorithm proceeds as follows (**Figure 3**):

1. Specify the parameter values that define the scenario (see *Section 4.7: Scenario-building and Sensitivity Testing*). Assuming the scenario under investigation is specific to OSW, specify the buildout goal (e.g., in terms of total GW of energy generated or total OSW development area) for the region(s). It is possible to include multiple regions with region-specific buildout goals in a single analysis by specifying additional constraints. Scenarios may have scenario-specific values for unknown parameters related to driving factors such as the effects of other anthropogenic activities on the receptor, climate change, current receptor abundance, life history parameters, or cause-effect pathways.
2. Specify the (likely) minimum size (area) of any single management area (for a regional assessment / region delineation CEA) or project (for a site selection CEA), *a_min_*.
3. Divide the area within the study area boundaries (e.g., for site selection, this refers to Newfoundland’s Preliminary Licencing Areas or Nova Scotia’s Potential Future Development Areas) into cells (*h*) that are smaller than *a_min_*. The algorithm can be constrained to select neighboring cells until the size of a given spatial cluster is at least *a_min_* (e.g., Ferguson et al. 2023). Allowing the algorithm to construct cells from smaller building blocks will result in the most effective use of space, assuming the species or pressure variables exhibit detectable spatial heterogeneity within *a_min_* and data with sufficient resolution are available. If the granularity of both the species and the pressure variables is larger than *a_min_*, then it is possible to obtain multiple solutions with different spatial configurations (i.e., different cells selected) but identical values of the CE metric.
4. Repeat the following steps for each iteration *i* until an optimal solution or a set of valid solutions is found:

a. Select a subset of cells, 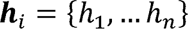 according to the pre-specified analytical constraints (including scenario parameter values).
b. Compute and save the CE metric 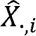 that is best suited to the information about the selected species and pressures. In the nomenclature for 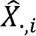, the dot specifies only one of the five CE metrics defined above. Below we address the more complicated scenario in which the available information varies across species or pressures, requiring computation of multiple types of CE metrics.
c. If the analytical objective is to find a single optimal solution, then:

i. If 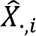 is the lowest value of the of the metric among all previous iterations, then save 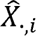 and ***h****_i_* as the new minimum value and associated solution, 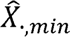 and ***h****_min_*, respectively.
d. If the analytical objective is to find a collection of valid solutions (i.e., collection of cells that meet all of the designated analytical constraints), then save ***h****_i_* and 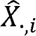 as a valid solution and its associated CE metric, respectively.

The spatial optimization analysis may be run using different parameters for scenario-specific parameters (*Section 4.7: Scenario-building and Sensitivity Testing*), *a_min_*, and cell size to examine the sensitivity of the results to these factors. Existing software such as Marxan (Ball, Possingham, and Watts 2009) or Marxan with Zones (Watts et al. 2009) are publicly available and free. Alternatively, customized computer code that applies linear integer programming (e.g., Ferguson et al. 2023) could be used and would provide the greatest flexibility for incorporating constraints specific to case studies; however, this approach would require additional time to create and debug the code.

For cases in which the available information varies across species or pressures, the spatial optimization analysis may be implemented independently for subsets of species or pressures using the relevant CE metrics. Recall from above that, for cases in which a given metric applies to multiple species, the contributions from each species are added together to compute the metric. Each implementation of the optimization algorithm would produce an optimal subset of sites, which would be solutions to the problem of minimizing potential CE of pressures on species based on the specific metric. In a subsequent step, the selected sites from each independent optimization analysis could be compared to identify where they do and do not overlap. Protocols should be specified in advance to weight the information from the different solutions in the following situations: cell *h* is present in all solutions; cell *h* is present only in the case for which a lot of information about the species or pressures was available; cell *h* is present only in the case for which relatively little information about the species or pressures was available; cell *h* contains individuals from a particularly vulnerable population (e.g., based on conservation status). A similar type of post hoc analysis could be used to combine results from species-focused CEAs with spatially explicit results from investigations focused on other aspects of the spatial planning issue, such as minimizing effects of OSW development on socioeconomic factors or on maximizing profit for the OSW industry. However, the results from this type of post hoc overlay analysis likely would differ from a comprehensive analysis that considers *all* constraints simultaneously.

The hierarchical spatial structure mentioned in Step 6 of *Section 3.4: Basic Steps for a Species-based CEA* is one way to incorporate the cumulative impacts of pressures that a species encounters outside of the OSW activity area. Based on knowledge or assumptions about the method in which multiple effects interact to impact a species (i.e., additive, synergistic, or countervailing), an additional step could be used to incorporate the effects of pressures encountered outside of the OSW activity area into the CE metric estimated using the spatial optimization algorithm. For example, if the species experiences heavy mortality due to bycatch in another portion of its range, then the estimated bycatch mortality could be added to an estimate of mortality produced using the total effect metric 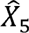. The total mortality could be used as input into a PVA.

## 6.0 Conclusions

As the offshore wind industry continues to expand, understanding and minimizing or mitigating its impacts on aerofauna is essential for sustainable development. This paper provides a comprehensive framework for conducting CEAs tailored to the early phases of planning and management of the OSW sector. Specifically, the framework can be used when the CEA is intended to inform regional assessment / region delineation or allocation of individual OSW project (licencing) areas inside predefined regional boundaries. The products developed using this framework may inform the site evaluation phase. For example, information about species and pressures that is compiled during the regional assessment / region delineation or site selection phases may be incorporated into future impact assessments.

Building on best practices, we introduce a species-centric approach to CEAs that can be used to guide the strategic siting of OSW developments to ensure they contribute to climate goals while protecting vulnerable aerofauna populations. Our analytical strategy enhances the flexibility and applicability of CEAs across various aerofauna receptor species or populations. The CE metric is defined as the product of a spatially explicit species variable and a spatially explicit pressure variable. We extended the utility of Halpern et al.’s (2008) approach by allowing the specific variables about receptors and pressures that are required to compute the CE metric to vary depending on the types of information that are available for the analysis. We explained how a number of superficially different approaches to conducting a CEA may be unified into a single generic analysis that may be applied to different information types, and we explain how analyses based on different information types can complement each other in a single analysis. We described spatial optimization methods that aim to minimize the value of the CE metric via the spatial allocation of individual polygons within a larger geographic area.

The ultimate advantages of this framework are that it can: (i) incorporate a broad range of spatially explicit information about species and pressures; (ii) allow for standardization of metrics and criteria across all assessments; and (iii) enable the results from different assessments to be compared, facilitating consistency in the decision-making process across space and time. While the framework was developed to support CEAs in Atlantic Canada, it is broadly applicable globally. Future research should focus on refining these methods through application to regional case studies.

## Acknowledgements

Thank you to the following experts who reviewed an earlier draft of this cumulative effects framework: Abel Gyimesi and Astrid Potiek (Waardenburg Ecology, The Netherlands), Kate Searle (UK Center for Ecology and Hydrology, United Kingdom), Edward Willsteed (Howell Marine Consulting Ltd, United Kingdom), Emma Kelsey (United States Geological Survey, USA), Noreen Kelly and Elizabeth Nagel (Fisheries and Oceans Canada, Canada).

## CRediT Statement

Megan C. Ferguson: Conceptualization, Methodology, Software, Writing - Original Draft, Writing - Review and Editing; Visualization, Project administration. **Kathryn A. Williams**: Conceptualization; Writing - Review and Editing, Supervision, Project administration. **M. Wing Goodale**: Writing - Review and Editing. **Evan M. Adams**: Writing - Review and Editing. **Paul Knaga:** Writing - Review and Editing. **Katrien Kingdon:** Writing - Review and Editing.**Stephanie Avery-Gomm:** Conceptualization; Writing - Review and Editing, Visualization, Supervision, Project administration, Funding acquisition.

## Footnotes

1 Nova Scotia: https://iaac-aeic.gc.ca/050/documents/p83514/147038E.pdf Newfoundland and Labrador: https://iaac-aeic.gc.ca/050/documents/p84343/147037E.pdf

2 https://cosewic.ca/index.php/en/status-reports.html

3 https://www.canada.ca/en/environment-climate-change/services/species-risk-public-registry/recovery-strategies.html

