## Appendix A for "A Flexible Framework for Assessing the Cumulative Effects of Offshore Wind Energy Activities and Other Pressures on Aerofauna"

### Appendix A: Glossary

**Action:** Any event that perturbs a receptor with a resultant effect (Masden et al. 2010).

**Additive effects:** If effects from multiple actions interact, and the total effect is the sum of its parts, then the effects are additive. If effects are not purely additive (e.g., they interact in ways that may increase or decrease the summed effect of the different actions), they may be termed synergistic, supra-additive, countervailing, or antagonistic effects (see below).

**Aerofauna:** Animals in the atmosphere, including birds, bats, and flying insects. For the purposes of this report, we use the term “aerofauna” to refer to only birds and bats.

**Baseline:** The initial, untouched, reference, or control state. In a CEA, the baseline sets the standard against which comparisons are made.

**Buildout:** The total extent of OSW that will be developed. It is typically measured by the total energy produced by OSW or the total area covered by wind farms. Buildout affects the magnitude or likelihood of a receptor species’ exposure to OSW activities.

**Countervailing/antagonistic effects:** If effects from multiple actions interact, and the total effect is less than the sum of its parts, then the effects are countervailing or antagonistic.

**Cumulative effects:** The combined effects of human activities and natural processes on wildlife across space and time (i.e., past, present, and reasonably foreseeable future).

**Cumulative effects assessment (CEA):** A systematic process of identifying, analyzing, and evaluating cumulative effects on a receptor ([CCME] Canadian Council of Ministers of the Environment 2014). Also referred to as “cumulative effects analysis”. The purpose of this systematic process is to inform planning and management.

**Cumulative effects metrics: T**he variable(s) or parameter(s) used to indicate the presence or magnitude of cumulative effects.

**Cumulative impacts assessment (CIA):** A systematic process of identifying, analyzing, and evaluating cumulative impacts on a receptor. Also referred to as “cumulative impacts analysis”. The purpose of this systematic process is to inform planning and management.

**Direct effect:** An effect on an action on a receptor.

**Effect:** The proximate response of an individual to an action (Masden et al. 2010).

**Endogenic pressures:** Pressures that emanate from within the system under study and can be managed or controlled (Elliott 2011; E. Willsteed et al. 2017).

**Exogenic pressures:** Pressures that emanate from outside the system or operate at scales beyond the system and cannot be directly controlled (Elliott 2011; E. Willsteed et al. 2017).

**Expert:** Someone who holds information about a given topic and who should be deferred to in its interpretation (Martin et al. 2012).

**Expert elicitation:** Structured expert elicitation is a tool that may be used to overcome concerns such as bias and lack of calibration when using expert judgment in decision-making. Martin et al. (2012) identify five steps in an expert elicitation: 1) deciding how information will be used; 2) determining what to elicit; 3) designing the elicitation process; 4) performing the elicitation; and 5) translating the elicited information into quantitative statements that can be used in a model or directly to make decisions.

**Expert judgment:** Predictions that arise when experts use their knowledge to predict what may happen in a particular context (Martin et al. 2012).

**Expert knowledge:** “[S]ubstantive information on a particular topic that is not widely known by others….This knowledge may be the result of training, research, and skills, but could also be the result of personal experience” (Martin et al. 2012).

**Exposure**: The frequency and duration by which individuals interact with a pressure over a specific geographic area (Goodale and Milman 2016).

**Impact:** The ultimate change in the fitness (reproduction or survival) of an individual or a population due to an action (Masden et al. 2010; Popper et al. 2022).

**Indirect effect:** An effect of an action on a receptor that is mediated by a third entity (Moon and Moon 2011).

**Licencing:** A verb describing the process of granting a license. Synonymous with permitting or granting.

**Observation uncertainty:** Observation uncertainty could be due to random errors (i.e., precision) or systematic discrepancies in magnitude or direction between data and reality (i.e., bias) (Stelzenmüller et al. 2020).

**Pathways of effects modelling**: A useful tool to systematically identify cause-effect pathways between anthropogenic activities, the associated stressors to ecological components, and the expected effects (Government of Canada 2012; Knights, Koss, and Robinson 2013).

**Planning phases:** Planning for offshore wind development can be divided into three phases: regional assessment and region delineation; site selection; and site evaluation. Regional assessment and region delineation describe the processes used to analyze and evaluate OSW development scenarios (e.g., defining the geographic boundaries within which OSW development may potentially occur), with the goal of informing and improving future planning, licencing, and impact assessment processes. Site selection occurs within pre-defined regional boundaries; it is the process of identifying specific boundaries for each project area that developers may bid on. Regional assessments, region delineation, and site selection encompass relatively large geographic areas. In contrast, site evaluation involves assessing the potential cumulative effects or impacts from a specific OSW project at a particular location, considered alongside a range of other OSW projects in the region or other types of activities influencing the same ecosystems and wildlife populations.

**Pressure scenario uncertainty:** Arises due to imperfect knowledge about the range and intensity of future human activities.

**Pressure scope:** Willsteed et al. (2018) define “pressure” in reference to CEAs as an “external abiotic or biotic factor exerted by an activity or other source that causes an effect”. The pressure scope is the set of pressures that are included in a CEA. Other terms used in the CEA field that refer to pressures with negative effects are “hazard” (Goodale and Milman 2016; Stelzenmüller et al. 2018), “impact-producing factors” (Bureau of Ocean Energy Management [BOEM] 2020; 2024), and “stressors” (Stelzenmüller et al. 2018). A single source may have multiple pressures. OSW is an example of a source; examples of pressures to birds from OSW include barrier to movement, lighting, and habitat alteration.

**Process uncertainty:** Process uncertainty in ecology refers to incomplete knowledge of relationships among natural phenomena, such as interactions among species and how species relate to their environment (Cressie et al. 2009). Ecological process uncertainty also includes incomplete knowledge of the relationships between natural phenomena and human activities (e.g., cause-https://www.canada.ca/en/impact-assessment-agency/services/policy-guidance/practitioners-guide-impact-assessment-act/guidance-describing-effects-characterizing-extent-significance.html pathways).

**Reasonably foreseeable:** The term “reasonably foreseeable” generally refers to something that is likely or expected to occur. Canada’s “Operational Policy Statement: Assessing Cumulative Environmental Effects under the Canadian Environmental Assessment Act, 2012”^[1]^ defines a “reasonably foreseeable physical activity” as one that is “expected to proceed, e.g. the proponent has publicly disclosed its intention to seek the necessary EA [Environmental Assessment] or other authorizations to proceed.” Similarly, the U.S. Code of Federal Regulations^[2]^ states that “reasonably foreseeable future actions include those federal and non-federal activities not yet undertaken, but sufficiently likely to occur, that a Responsible Official of ordinary prudence would take such activities into account in reaching a decision. These federal and non-federal activities that must be taken into account in the analysis of cumulative impact include, but are not limited to, activities for which there are existing decisions, funding, or proposals identified by the bureau. Reasonably foreseeable future actions do not include those actions that are highly speculative or indefinite.”

**Receptor:** A feature that is sensitive to, or has the potential to be affected by, an action (Masden et al. 2010).

**Regional assessment:** Under the authority of the *Impact Assessment Act* (2019), the Nova Scotia and the Newfoundland and Labrador Regional Assessment Committees are tasked with identifying and considering the potential positive and adverse effects, including cumulative effects, of future offshore wind development activities in the regional Study Areas, as well as “potential interactions between the effects of future offshore wind development activities and those of other existing and future physical activities, including the potential for resulting cumulative effects” (Committee for the Regional Assessment of Offshore Wind Development in Nova Scotia 2024; Committee for the Regional Assessment of Offshore Wind Development in Newfoundland and Labrador 2024).

**Scope**: The envelope or boundaries defining the sources, pressures, space, time, and receptors that constrain an analysis.

**Sensitivity**: Gușatu et al. (2021) define sensitivity as “the likelihood of change when a pressure is applied to a receptor (environmental component) and is a function of the ability of the receptor to adapt, tolerate or resist change and its ability to recover from the impact”.

**Source scope**: The anthropogenic activities or environmental drivers whose actions may affect the receptors and that are included in the CEA. A single source may have multiple pressures. OSW is an example of a source; examples of pressures to birds from OSW include barrier to movement, lighting, and habitat alteration.

**Spatial scope**: The spatial extent of the analysis.

**Synergistic/supra-additive effects**: If effects from multiple actions interact, and the total effect is greater than the sum of its parts, then the effects are synergistic or supra-additive.

**Statistical uncertainty**: Uncertainty in the assumptions used in statistical models; examples include uncertainty in model structure or parameter estimates (Stelzenmüller et al. 2020).

**Temporal scope**: The period (seasons and years) included in the analysis.

**Uncertainty**: The state of deficiency of information related to understanding or knowledge of an event, its consequences or likelihood (Stelzenmüller et al. 2018).

**Valued Component (VCs)**: Elements of the human and natural environment that are important to participants in an impact assessment process. Valued components are identified by Indigenous communities, the public, federal authorities or proponents. They may have scientific, biological, social, cultural, economic, historical, archaeological or aesthetic importance, and may be intricately related to community health and well-being (IAAC, 2023).

**Vulnerability**: Vulnerability is a function of exposure and sensitivity. Goodale and Milman (2016) define it as “the likelihood an individual will interact with, and respond to” a pressure, “and that the response will adversely affect the population”.
